# Early eye and forebrain development are facilitated by Bone Morphogenetic Protein antagonism

**DOI:** 10.1101/2022.12.01.518698

**Authors:** Johannes Bulk, Valentyn Kyrychenko, Philipp Rensinghoff, Stephan Heermann

## Abstract

Vision likely is our most prominent sense and a correct development of the eye is at its basis. Early eye development is tightly connected to the development of the forebrain. A single eye field and the prospective telencephalon are situated within the anterior neural plate (ANP). If development is running correctly both are split and consecutively two optic vesicles and two telencephalic lobes emerge. If hampered, the domain is remaining condensed at the midline. This affection of development is termed Holoprosencephaly (HPE). The classical ocular finding associated with intense forms of HPE is cyclopia, one central eye.

We found that antagonists of Bone morphogenetic proteins (BMP) are important to facilitate proper forebrain and eye field cleavage. Experimental induction of a BMP ligand results in HPE and the analyses of the ANP indicated a severe form. We further found anophthalmia instead of cyclopia associated with the present HPE phenotype. We identified retinal progenitors stuck in the forebrain domain, which we termed crypt-oculoid. Our data further suggest that the process of basal constriction of retinal progenitors is hampered by elevated levels of the BMP ligand. This likely is the reason for anophthalmia instead of cyclopia in this present case of HPE.

## Introduction

Vision likely is our most prominent sense. A correct development of the light sensing organ, the eye, is at its basis. Overall, eye development is an intricate yet fascinating process. During gastrulation a single eye field within the anterior neural plate (ANP) is divided and subsequently two optic vesicles emerge at the lateral surface of the anterior neuroectoderm. The progenitors within the eye field behave like single cells and also differently compared to the telencephalic progenitors. During neurulation the telencephalic progenitors converge towards the midline while the eye field progenitors stay more laterally and then move even further away from the midline (Rembold et al., 2006). This differential behavior is supported by a *rx3* driven suppression of *alcamb* in the eye field progenitors and an active expression of *alcamb* in the *rx3* negative telencephalic progenitors (Brown et al., 2010). Importantly, the division of the ANP is also supported by a subduction movement of hypothalamic progenitors. These progenitors are located posterior to the eye field originally but then migrate ventrally and rostrally inducing neural keel formation. This movement somehow indirectly supports the division of the ANP (England et al., 2006). But not only the eye field has to be divided, also the prospective telencephalic domain within the ANP has to be split, eventually resulting in two telencephalic lobes. Failures during these processes cause holoprosencephaly (HPE). The spectrum of HPE phenotypes varies between mild forms and intense forms containing the classical ocular phenotype being cyclopia, but also anophthalmia and coloboma (Fallet-Bianco, 2018). HPE is mostly genetically linked and the most prominent factor important for proper ANP division is Sonic hedgehog (Shh), derived from the prechordal plate (Chiang et al., 1996; Roessler et al., 1996).

BMP signaling must be activated at a specific level during the initiation of the ANP to facilitate the expression of Wnt antagonists in the anterior neural border (Houart et al., 2002) which in combination with Wnt ligand expression in posterior domains are ensuring a functional Wnt gradient over the ANP (Cavodeassi, 2014). This is important for ANP domain specification. Slightly later, BMP signaling was found to be favoring the telencephalic progenitor fate over the fate of eye field progenitors (Bielen and Houart, 2012). BMP signaling can be affected by different means, e.g. by regulation of ligand or receptor expression but also by induction of BMP antagonists. BMP antagonists, derived from the mouse node and axial mesendoderm, were found to be essential for head development. A combined loss of chordin and noggin affected the formation of the prosencephalon and also resulted in cases of cyclopia (Bachiller et al., 2000), thus providing embryos with variable intense forms the HPE spectrum. Notably, such a loss of BMP antagonists could result in an overactivation of BMP signaling. Yet, an enlargement of the telencephalon at the expense of the eye field, as seen after BMP signaling activation in zebrafish (Bielen and Houart, 2012), was not described resulting from the loss of BMP antagonists in mouse. This may be due to species differences (Bachiller et al., 2000).

In the recent years it became evident that also the consecutive step of eye morphogenesis, especially the transformation from the optic vesicle to the optic cup, is a highly dynamic process (Heermann et al., 2015; Kwan et al., 2012; Picker et al., 2009; Sidhaye and Norden, 2017). Besides, the driver for neuroretinal bending during optic cup morphogenesis was found, *opo/ofcc1* (Martinez-Morales et al., 2009a). Ofcc1 is supporting basal constriction of individual neuroretinal precursors and thus the drastic shape change of the entire developing optic cup (Bogdanović et al., 2012; Martinez-Morales et al., 2009a). Further, it could be shown that the morphogenetic movements and the *ofcc1* function are linked (Sidhaye and Norden, 2017). The dynamic cell movements which bring lens averted cells into the lens facing domain and which are also important for optic fissure morphogenesis are further dependent on BMP antagonism (Eckert et al., 2019; Heermann et al., 2015).

Experimental induction of a BMP ligand results in a precocious arrest of these movements, ectopic retina and in a specific form of coloboma, a “morphogenetic coloboma” (Eckert et al., 2019; Heermann et al., 2015; Knickmeyer et al., 2018). We hypothesize that such a morphogenetic coloboma is more likely linked to HPE than a coloboma resulting from a fusion defect of the optic fissure margins and an affected pioneer cell population (Chan et al., 2021; Eckert et al., 2020). We thus propose that BMP antagonism is involved also in early forebrain development.

## In this study

we addressed the role of BMP antagonism during ANP and eye field development. BMP antagonists are frequently found expressed redundantly, underlining their pivotal role during embryogenesis (Bachiller et al., 2000; Khokha et al., 2005). We found BMP antagonists expressed in the ANP at 11 hpf. To saturate this domain, we experimentally induced BMP expression at 8.5 hpf. This induction resulted in anophthalmia with *rx2* positive retinal progenitor cells being stuck in the forebrain. Analysis of ANP domains at 11 hpf showed a failure of telencephalic field and eye field division and a loss of *rx3* expression, while *shha/b* expression was strongly reduced, but not completely absent. Analysis of emx3 expression at 24 hpf points at least to a partial failure of the hypothalamic subduction movement leaving a part of the emx3 domain posterior to the eye anlagen. Notably, even though retinal precursors were clearly identified inside the brain, no eye, not even a single cyclopic eye, was formed. A delayed induction of BMP, allowing the optic vesicle evagination to begin, resulted in a flat surface of the optic vesicle and prevented optic cup and eye formation. We propose that the induction of BMP negatively affects *ofcc1* function and thus proper optic cup and eye formation. This could be the reason why also the retinal progenitors, situated close to the midline are not transformed into a cyclopic eye.

## Results

### *bmp4* induction results in HPE and a “crypt-oculoid”

HPE has various possible phenotypic manifestations and coloboma can be one. Previously we described the role of BMP antagonism for the transformation of the optic vesicle to the optic cup (Eckert et al., 2019; Heermann et al., 2015) and its implication for the formation of a morphogenetic coloboma. We hypothesize that such morphogenetic coloboma are potentially linked to HPE. Here, we investigated whether BMP antagonists could also be important for division of the ANP including the eye field. We found BMP antagonists, *grem2b* and *fsta*, expressed in the and neighboring to the ANP respectively at 10, 11 and 12 hpf (**Figure 1 A-M, dotted lines and arrows**). In order to oversaturate this domain containing the BMP antagonists and to potentially affect ANP development, we used a transgenic line allowing for a heat shock induced expression of *bmp4* (*tg(hsp70l:bmp4, cmlc2:GFP)*) and consecutive signaling activation (Eckert et al., 2019; Eckert et al., 2020; Knickmeyer et al., 2018) and performed *bmp4* induction at 8.5 hpf (**Figure 1 N**). First, we performed a gross morphological analysis at 48 hpf with a stereomicroscope. At this age, the embryos harboring the transgene could be identified by the transgenesis marker (*cmlc2:GFP*).

**Figure 1.**
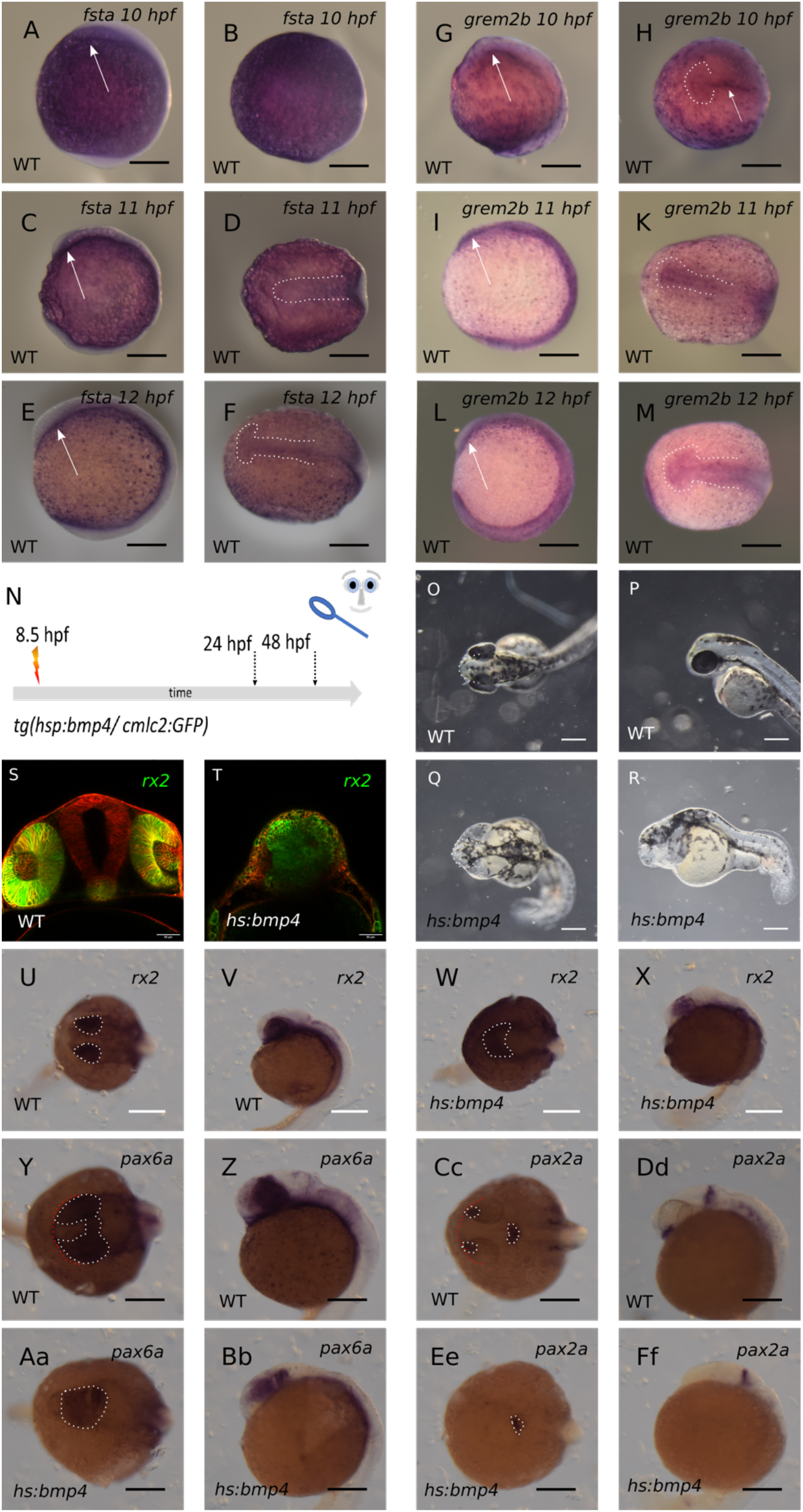
**A-M: expression of bmp-antagonists *fsta* and *grem2b* in the ANP at 10, 11 and 12 hpf. A-F: ISH for *fsta*. A, B: 10 hpf; C, D: 11 hpf; E, F: 12 hpf. G-M: ISH for *grem2b*. G, H: 10 hpf; I, K: 11 hpf; L, M: 12 hpf. A, C, E, G, I, L: lateral view. B, D, F, H, K, M: dorsal view. Scalebars indicate 200 µm.** **N: summary of experimental procedure. Embryos were heat shocked at 8,5 hpf and observed at 24 hpf or 48 hpf, resp.** **O, P: bright-field image of wild-type at 48 hpf. Q, R: bmp4-induced larva at 48 hpf. O, Q: dorsal view. P, R: lateral view. Scalebars indicate 250 µm.** **S, T: transversal confocal sections of larvae with rx2:GFP at 24 hpf. injection of *LY-tdTomato* mRNA in zygote. S: wild-type. T: bmp4-induced, scalebars indicate 50 µm.** **U-Ff: ISH for *rx2* (U-X), *pax6a* (Y-Bb) and *pax2a* (Cc-Ff) at 24 hpf. U, V, Y, Z, Cc, Dd: wild-types. W, X, Aa, Bb, Ee, Ff: bmp4-induced. U, W, Y, Aa, Cc, Ee: dorsal view; V, X, Z, Bb, Dd, Ff: lateral view. scalebars indicate 250 µm (U-X) or 200 µm (Y-Ff) resp.** **Red lines indicate outlines of heads, dotted lines indicate the expression domains.**

Embryos without the transgene served as controls (**Figure 1 O, P**). Following the *bmp4* induction, we found a severe defect in the developing forebrain and could not detect any eyes (**Figure 1 Q,R**). We next asked whether residual retinal tissue could be existent. To test this, we added another transgenic zebrafish line *tg(rx2:GFPcaax)* to our analysis in which retinal progenitors express a fluorescent protein and used it in combination with the *tg(hsp70l:bmp4, cmlc2:GFP)* and induced *bmp4* expression at 8.5 hpf. Surprisingly, using confocal microscopy for analysis, we found many GFP expressing cells inside the forebrain, closer to the midline, compared to the control, yet no eye was formed (**Figure 1 S, T**).

Besides the GFP, driven by the *rx2* cis regulatory element, we also found the *rx2* transcript with whole mount in situ hybridization (WMISH) (**Figure 1 U-X, dotted lines**). Also, the expression of the transcript clearly showed that retinal progenitors are present and that they are localized closer to the midline, compared to controls, like a “crypt-oculoid” (**Figure 1 S, W, X**). We further performed WMISH for *pax6a* and *pax2a*, two important transcription factors for eye development and axis specification. *Pax2* is important for proximal fates and optic stalk development and *pax6* for optic vesicle/cup development (Chow and Lang, 2001) (**Figure 1 Y-Ff**). While in controls we found *pax6a* expressed in the optic cups and diencephalon, we found it expressed in the domain of the crypt-oculoid and a domain posterior to this, likely corresponding to the diencephalic domain, after *bmp4* induction (**Figure 1 Y-Bb, dotted lines**). *Pax2a* in controls on the other hand marked the optic stalk and the midbrain hindbrain boundary (**Figure1 Cc, Dd, dotted lines**). After *bmp4* induction, the midbrain hindbrain boundary was still visibly expressing *pax2a*, while optic stalks could not be identified (**dotted line Figure 1 Ee, Ff**). Together, this indicates that the induction of *bmp4* at 8.5 hpf resulted in a failure of eye field separation and optic vesicle out-pocketing. It also showed that even though retinal precursors, positive for *rx2* and *pax6a*, were present, no eye was formed.

### ANP development is hampered after *bmp4* induction

Having seen that *bmp4* induction results in a failure of eye field splitting, we next more closely addressed the effect on the development of the ANP. To this end, we fixed embryos at 11 hpf and processed the embryos for WMISH (**Figure 2 A**). We used markers for the prospective forebrain (*emx3* and *foxg1*) domain and the prospective eye field (*six3b* and *rx3*). The transgenesis marker (*cmlc2:GFP*) was not yet active at 11 hpf. Thus, the WMISH was performed blinded and the genotype (wt-controls vs. *hspl:bmp4*) was determined after the WMISH and the documentation of the results. It could be expected that an excess of a BMP ligand is potentially resulting in an expansion of the prospective telencephalic domain at the expense of the eye field (Bielen and Houart, 2012). But we could not detect a shift in fate in between these two domains. We rather found that resulting from the induction of *bmp4*, the forebrain domain (*emx3*) and the eye field (*six3b*) were both condensed at the midline (**Figure 2 B-E dotted lines and arrows, F-I**). This indicates that the induction of *bmp4* was hampering the division of the ANP containing both the eye field and the prospective forebrain domain, being the hallmark of HPE. Interestingly, the expression of *rx3* and *foxg1a* were ceased/lost after *bmp4* induction (**Figure 2 K-R, dotted line and arrows**). This suggests that either the induction or the maintenance of the expression of these two transcription factors was sensitive to elevated levels of *bmp4. Foxg1* is an essential transcription factor involved in many aspects of telencephalic development including also the growth of this brain region (Hettige and Ernst, 2019). *Foxg1* was also shown to be essential for the development of the olfactory system including the olfactory epithelium and also the olfactory bulb, a telencephalic region (Duggan et al., 2008). A loss of *foxg1* expression could thus likely be resulting in reduced size of the telencephalon. A loss of *rx3* could be a plausible reason for the anophthalmia phenotype (Bielen and Houart, 2012; Brown et al., 2010; Loosli et al., 2003; Rembold et al., 2006). It was shown that *rx3* was important for maintenance of the chemokine receptor *cxcr4* within the eye field (Bielen and Houart, 2012), which in combination with *sdf1b* from the underlying mesendoderm is supposed to be important for eye field evagination. We found a strongly reduced level of *cxcr4* expression in the eye field after induction of *bmp4* (**Figure 2 S-V**), in line with the previous findings (Bielen and Houart, 2012) and supporting the idea that the loss of *rx3* and consecutively loss of cxcr4 is the reason for anophthalmia after induction of *bmp4*. Furthermore, it was shown with previous data that a loss of *rx3* was resulting in an upregulation of *alcamb* (*nlcam*) within the eye field progenitors (Brown et al., 2010) making them behave like their telencephalic neighbors. It was unexpected to see that in controls at 11 hpf the expression domain of alcamb was not yet corresponding and restricted to the telencephalic domain (**Figure 2 W dotted line, X**), as demonstrated for a 6 somite staged embryo at 12 hpf (Brown et al., 2010). After induction of *bmp4* we found an overall reduced level of *alcamb* within the ANP (**Figure 2 Y dotted line, Z**).

**Figure 2.**
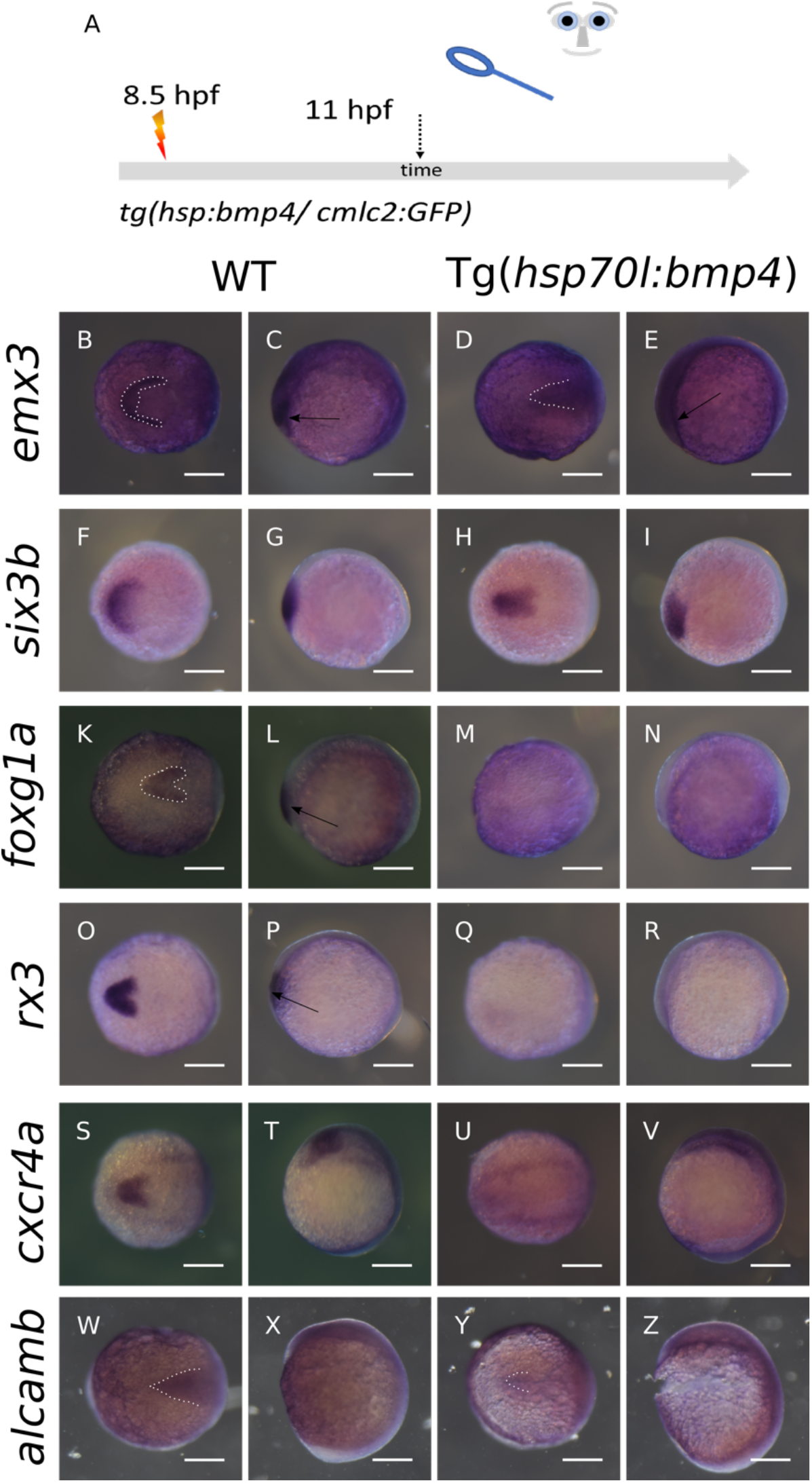
**ISH at 11 hpf after bmp4-induction at 8,5 hpf. left columns wild-types, right columns bmp4-induced. first and third column dorsal view, second and fourth column lateral view. A: summary of experimental procedure. Embryos were heat-shocked at 8,5 hpf and analyzed at 11 hpf. B-E: Expression of *emx3* is condensed at the midline and weakened (dotted lines and arrows). F-I: expression of *six3b* is also condensed at the midline. K-N: *foxg1a* (dotted line and arrow) is lost after bmp4-induction. O-R: *rx3* (arrow) is lost as well. S-V: *cxcr4a* is lost. W-Z: expression of *alcamb* (dotted lines) is weakened. Scalebars indicate 200 µm.**

We found it surprising that *rx3* was lost after *bmp4* induction while *rx2* positive progenitors could still be detected at a later stage of development (**Figure 1 U-X /2 O-R**). It was shown in the zebrafish *rx3* mutant *chokh* that *rx2* expression is depending on *rx3* expression (Loosli et al., 2003). In Medaka, however, it was shown that a loss of *rx3* is not resulting in a loss of *rx2* expression (Loosli et al., 2001). We next targeted the *rx3* locus of the zebrafish genome with CRISPR/Cas9 and analyzed F0 Crispants (**Figure 3 A**), technically similar to previous analyses of Wu and colleagues (Wu et al., 2018). We reproducibly observed anophthalmia and severe microphthalmia in our F0 Crispants (**Figure 3 B-E**). We next performed WMISH of F0 *rx3* Crispants using an *rx2* probe (**Figure 3 F-I**). Notably, we found highly reduced to absent expression levels of *rx2* in our F0 *rx3* Crispants, correlating nicely with the efficacy of the CRISPR induced anophthalmia phenotype and supporting the idea that *rx2* is dependent on *rx3* expression in zebrafish. However, this finding is at odds with our observation, that *rx3* is absent after *bmp4* induction, while rx2 is present afterwards (**Figure 1 W, X/2 Q, R**).

**Figure 3.**
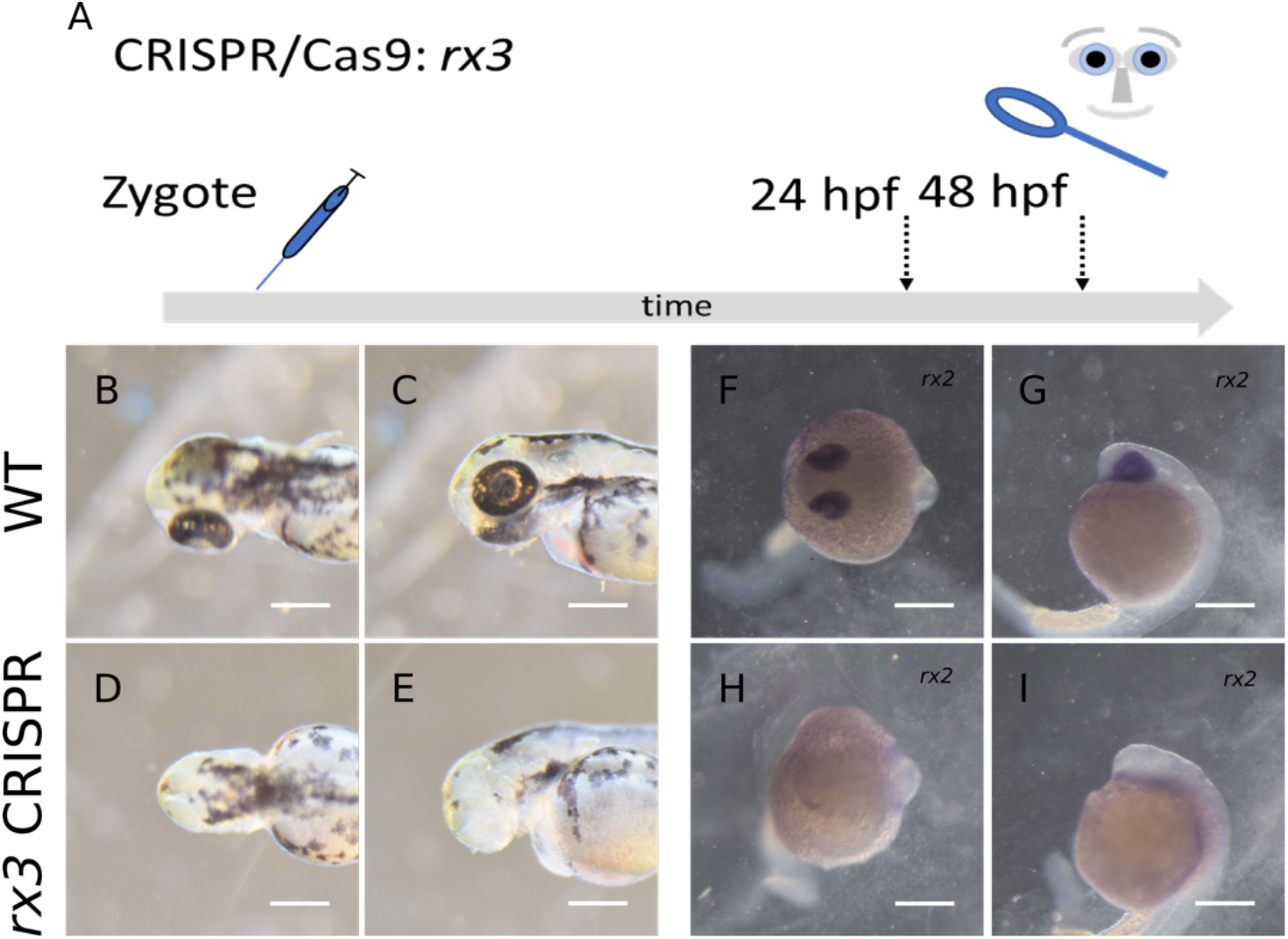
**A: summary of experimental procedure: embryos were injected at 1-cell-stage and analyzed at 24 hpf or 48 hpf. B-E: Brightfield-images at 48 hpf. F-I: ISH for *rx2* at 24 hpf. B, C, F, G: wild-type. D, E, H, I: *rx3*-crispant. B, D, F, H dorsal view. C, E, G, I lateral view. scalebars indicate 250 µm.**

### *Zic2a* expression ceased and *shha* is reduced in the prechordal plate after *bmp4* induction

*Bmp4* induction results in HPE. The phenotype was showing a splitting defect of the ANP involving the prospective forebrain and the eye field. The most prominent reason underlying HPE is the loss of Shh secreted from the prechordal plate. Next, we addressed the expression of *shha/b* at 11 hpf in controls and *bmp4* induced embryos (**Figure 4 A-I**). In controls, we found *shha* and *shhb* expressed in the axial mesendoderm (**Figure 4 B, C, F, G**). The expression of *shha/b* was reaching far to the anterior end of the embryo, where the domain was broadened and showing strong expression (**arrows in Figure 4 C, G**) corresponding to the prechordal plate. After induction of *bmp4*, we found the expression of both *shha* and *shhb* variably reduced (**Figure 4 D, E, H, I**), yet, not absent. Notably, the anterior domain of *shha* was less broad and less intense, suggesting an affected prechordal plate (**Figure 4 D, E dotted line and arrows**). The transcription factor *zic2* also is an established HPE related gene. If mutated it hampers prechordal plate development (Warr et al., 2008) and thus results in HPE. Later in development, *zic2* was, however, also suggested to act downstream of Shh, limiting the expression of *six3* in the developing forebrain (Sanek et al., 2009). We addressed *zic2* expression after *bmp4* induction (**Figure 4 K-N**). While in controls *zic2* is expressed among other domains at the border of the ANP, we found *zic2* almost absent in this region after *bmp4* induction (**Figure 4 M, N**). In our paradigm we perform *bmp4* induction, to saturate domains of BMP antagonists in the ANP (**Figure 1 A-N**). After induction of *bmp4*, we find reduced expression of *zic2* in the border of the ANP and an altered domain of especially *shha* expression in the prechordal plate region. Together our findings suggest, that BMP antagonists are important for *zic2* induction or maintenance in the ANP region and proper establishment of the prechordal plate to eventually facilitate proper division of the telencephalic domain and the eye field. Whether the reduction of *zic2* is resulting in a defective prechordal plate, whether zic2 is (in)active downstream of shh or whether both are affected independently is yet unsettled.

### *Bmp4* induction likely affects neural keel formation and hypothalamic subduction

While *emx3* is a marker for the prospective telencephalon at early stages (11 hpf), it is also expressed within the diencephalon/hypothalamus at later embryonic stages (Jung et al., 2020; Paridaen et al., 2009). We addressed the expression of emx3 at 24 hpf (**Figure 5 A-E**). In controls *emx3* is mainly showing expression in the developing telencephalon and mild expression in the ventral diencephalon both in anterior parts, related to the eye anlagen (**Figure 5 B, C dotted line and arrows)**. After *bmp4* induction, *emx3* is broadly expressed caudal to the residual eye anlage. Compared to the control, a smaller *emx3* positive domain is visible in the residual telencephalic region also entangling the crypt oculoid (**Figure 5 D, E dotted line and arrows**). As mentioned above, the reduction in size of the telencephalic *emx3* positive domain can be explained by the loss of *foxg1* expression in the ANP (**Figure 2 K-N**). The localization of the domain posterior to the eye anlage suggests at least a partial failure of the hypothalamic subduction movement.

**Figure 4.**
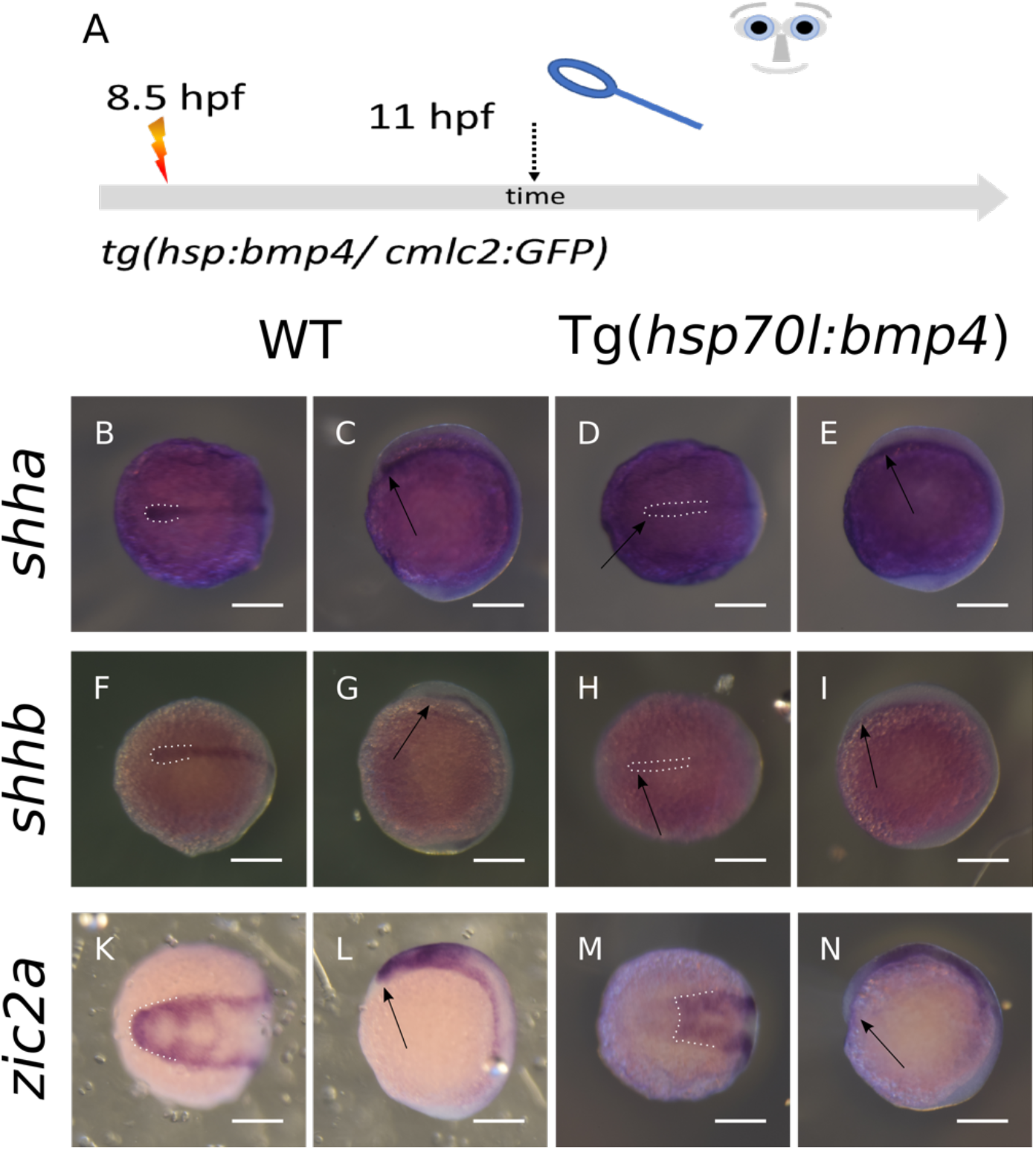
**ISH at 11 hpf after bmp4-induction at 8,5 hpf. A: summary of experimental procedure: embryos were heat-shocked at 8,5 hpf and analyzed at 11 hpf. left columns wild-types, right columns bmp4-induced. first and third column dorsal view, second and fourth column lateral view. B-E: Expression of *shha* is reduced after *bmp4*-induction yet still present (dotted lines and arrows). F-I: *shhb* is also still present but reduced in intensity (dotted lines and arrows). K-N: Expression of *zic2a* is lost in the ANP domain after *bmp4*-induction (dotted lines and arrows). scalebars indicate 200 µm.**

**Figure 5.**
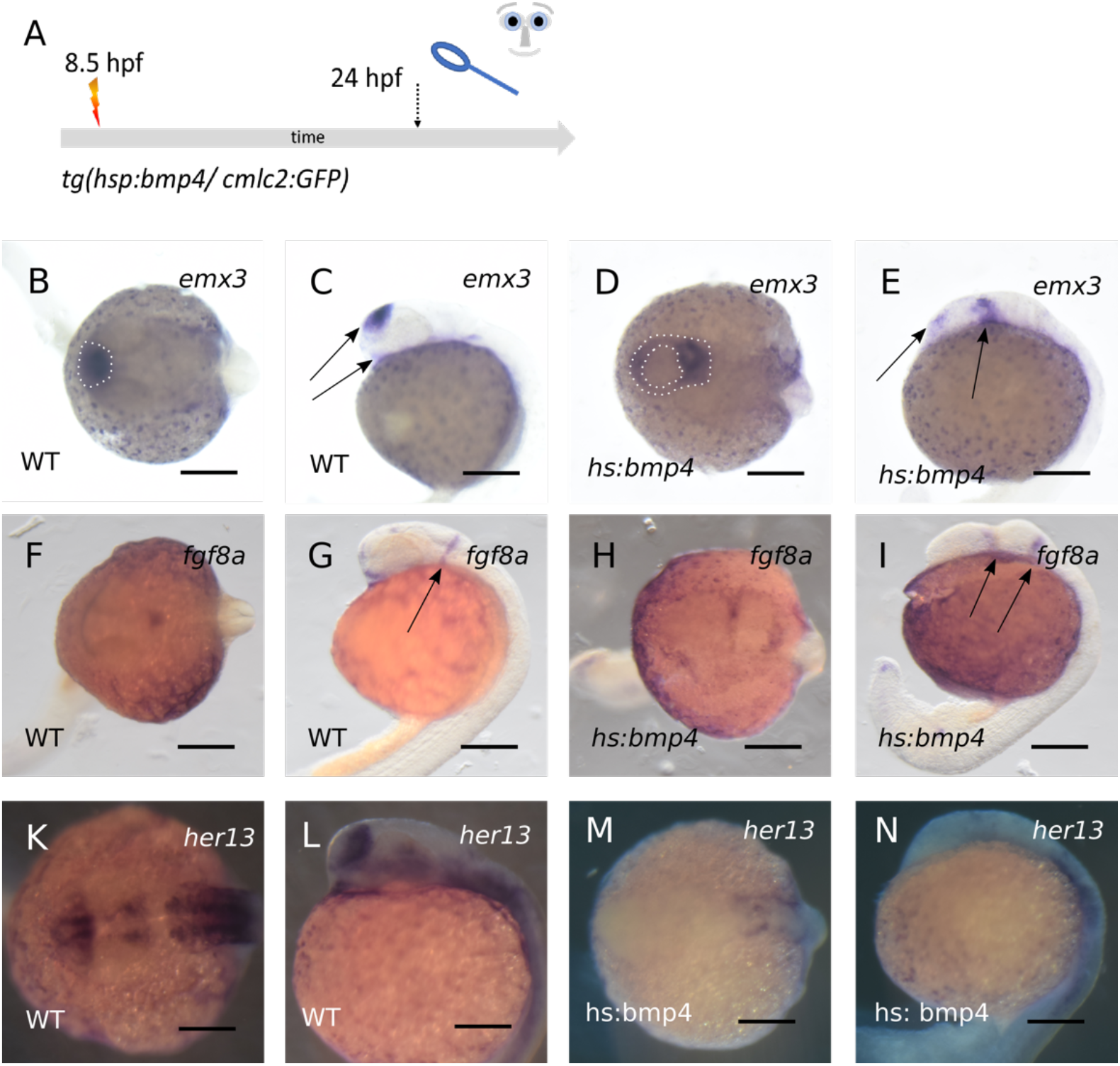
**A: summary of experimental procedure: embryos were heat-shocked at 8,5 hpf and analyzed at 24 hpf. B-E: ISH for *emx3* at 24 hpf. *Emx3* is present after bmp4-induction but the expression pattern is changed (arrows). F-I: ISH for *fgf8a* at 24 hpf. The *fgf8a* domain in the forebrain has changed after *bmp4*-induction (arrows). K-N: ISH for *her13* at 24 hpf. The expression of *her13* is lost in anterior regions after bmp4 induction. B, C, F, G, K, L: wild-types. D, E, H, I M, N: *bmp4*-induced. B, D, F, H, K, M: dorsal view; C, E, G, I, L, N: lateral view. scalebars indicate 200 µm.**

The subduction of the hypothalamic precursors during normal development is important to establish the neural keel in the anterior region and thus extend the dorsal ventral axis (England et al., 2006). The induction of *bmp4*, however, is resulting in a more linear arrangement of the telencephalic domain, the eye field and the hypothalamic domain, from anterior to posterior. As beforementioned, we found a broad emx3 expressing domain posterior to the crypt-oculoid. It cannot be excluded, that also telencephalic precursors are misplaced here (**Figure 5 E right arrow**).

We next addressed the expression of other genes marking in part diencephalic identity. We addressed the expression of *fgf8a* (**Figure 5 F-I**). In controls *fgf8a* is expressed in the midbrain hindbrain boundary and in anterior regions, including the ventral diencephalon (**Figure 5 F, G arrow**). After induction of *bmp4* the anterior expression is ceased, while two expression domains posterior to the oculoid could be detected (**Figure 5 H, I arrows**). We also addressed the expression of *her13*, which in controls is expressed in diencephalic and telencephalic regions (**Figure 5K, L**). After *bmp4* induction, however, the expression of *her13* is largely lost, especially in anterior regions (**Figure 5 M, N**). While the loss of *her13* expression suggest a defect in growth and differentiation, the finding of the two *fgf8a* domains posterior to the oculoid is in line with a defect of subduction and forward migration.

### Anophthalmia instead of cyclopia results from *bmp4* induction

We identified a form of HPE showing an ANP splitting defect resulting from an induction of *bmp4*. We also identified eye field precursors inside the forebrain, but unexpectedly, instead of a cyclopic condition we found anophthalmia. We thus asked next, why is anophthalmia resulting from *bmp4* induction instead of cyclopia? It is conceivable or at least possible that anophthalmia is an even more severe phenotype than cyclopia?

We next addressed whether a milder induction of *bmp4* would result in HPE with cyclopia instead of anophthalmia. To this end we used less intense heat shock paradigms for induction of *bmp4*, 7.5 minutes and 10 minutes instead of 15 minutes (**Figure 6 A**). Both milder induction paradigms resulted in milder forms of HPE (**Figure 6 B-G**). In these embryos, however, two eye analgen were formed respectively, with morphological aberrations and, importantly, a smaller distance between the eyes, ocular hypotelorism, a phenotype of milder HPE (**Figure 6 D, G**). We could not detect cyclopic embryos. This showed that the level of bmp4 was important for the intensity of HPE and that in case of HPE induced via *bmp4* induction, a strong form shows anophthalmia and milder forms show ocular hypotelorism compared to the wild-type (**Figure 6 H, I**). We hypothesize that cyclopia is not milder but rather different than anophthalmia. We next addressed further, why anophthalmia is resulting from *bmp4* induction.

**Figure 6.**
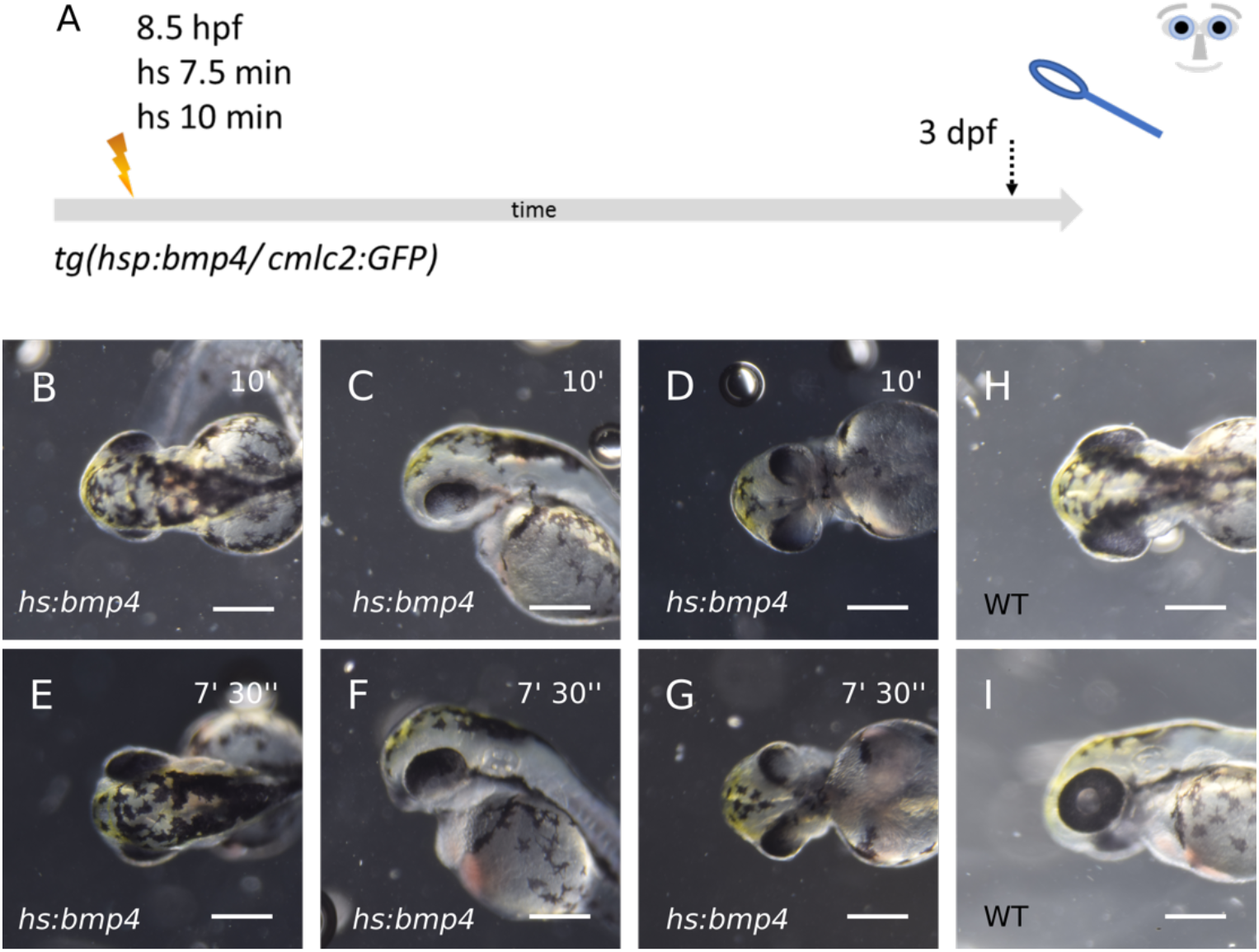
**Inefficient bmp4-induction does not lead to cyclopia. A: summary of experimental procedure: embryos were heat-shocked at 8,5 hpf for 7,5 or 10 min and analyzed 3 dpf. : *bmp4*-induction for 10 min instead of 15 min. The fish develop two optic cups that are fused in the middle. E-G: *bmp4*-induction for 7,5 min. The fish develop a wide coloboma. H, I: Wild-types at 3 dpf. B, E, H: dorsal views, C, F, I: lateral views, D, G: ventral views. Scalebars indicate 250 µm.**

An interaction between the pre-lens ectoderm and the optic vesicle is fundamental for the transformation of the optic vesicle into the optic cup (Hyer et al., 2003). A lens could not be identified by gross morphological analysis (**Figure 1 Q, R**) after *bmp4* induction. We next asked if nonetheless lens tissue was detectable after a strong induction of *bmp4* at 8.5 hpf (**Figure 7 A)**. We addressed the expression of *cryaa* by WMISH (**Figure 7 B-E)**. *Cryaa* is nicely expressed in the control lenses (**Figure 7 B, C**). Even though no lens could be identified, we detected *cryaa* expression anterior to the crypt-oculoid, in the region where a lens would be formed in a cyclopic eye (**Figure 6 D, E**). Pre-lens ectoderm and retinal progenitors are both present, yet no eye is formed after induction of *bmp4*. In order to be transformed into an optic cup, the individual retinal progenitors must undergo a change of shape. They turn from a columnar into a wedge form by basal constriction, which is facilitated by the protein Opo/Ofcc1 (Bogdanović et al., 2012; Martinez-Morales et al., 2009b). We next addressed whether a strong induction of *bmp4* (15 minutes), however, performed at a stage when the optic vesicle out-pocketing has been initiated already, would affect optic cup formation. To this end we induced *bmp4* at 10.5 hpf instead of 8.5 hpf (**Figure 7 A**). We used the transgenic line *tg(rx2:GFPcaax)* in order to visualize the retinal progenitors. While in controls the optic cup formation was visible nicely, the delayed but strong *bmp4* induction resulted in optic vesicles with a rather flattened surface (**Figure 7 F, G**) and lens tissue was not found (Figure 6 H-I). The flat surface of the optic vesicle was reminding of *opo* mutants (Martinez-Morales et al., 2009b) and was clearly indicating a hampered optic cup formation.

**Figure 7.**
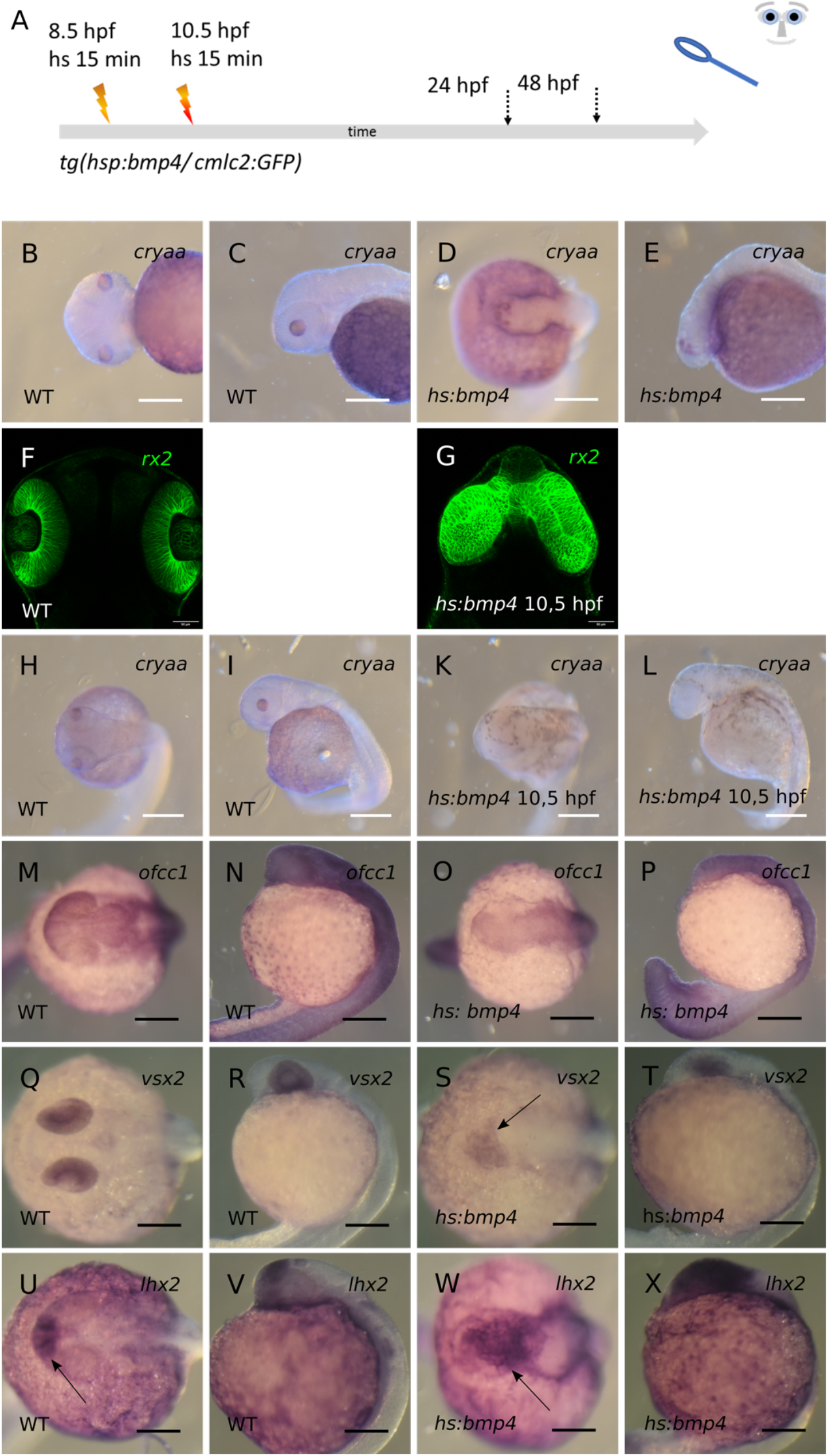
**A: summary of experimental procedure: embryos were heat-shocked at 8,5 hpf or 10,5 hpf for 15 min and analyzed 24 or 48 hpf. : ISH for *cryaa* at 48 hpf after a 15 min heat-shock at 8,5 hpf. B, D: dorsal views. C, E: lateral views. B, C: wild-type. D, E: *bmp4*-induction for 15 min at 8,5 hpf. The embryo has no eyes and multiple small *cryaa*-domains at the tip of the head. Scalebars indicate 200 µm.** **F, G: Confocal images with green indicating *rx2*-positive tissue at 24 hpf. F: wild-type. G: bmp4-induction at 10,5 hpf. Two optic vesicles without a lens have formed. Coronary sections. Scalebars indicate 50 µm.** **H-L: ISH for *cryaa* at 48 hpf after a 15 min heat-shock at 10,5 hpf. H, I: wild-type. K, L: *bmp4*-induction for 15 min at 10,5 hpf. The embryo has no eyes and no visible *cryaa-* positive tissue. H, K: dorsal view. I, L: lateral view. Scalebars indicate 250 µm.** **M-X: ISH at 24 hpf after 15 min heat-shock at 8,5 hpf. First and second columns wild-types; third and fourth columns *bmp4*-induced. First and third columns dorsal views, second and fourth columns lateral views. M-P: ISH for *ofcc1. Ofcc1* is still present in the oculoid after bmp4-induction. Q-T: ISH for *vsx2. Vsx2* is expressed in the optic cup of wild-types and in the oculoid (arrow). U-X: ISH for *lhx2. Lhx2* is also expressed in the oculoid after bmp4-induction as well as in the telencephalon, diencephalon and ventral optic cup in wild-types (arrows). Scalebars indicate 200 µm.**

**Figure 8.**
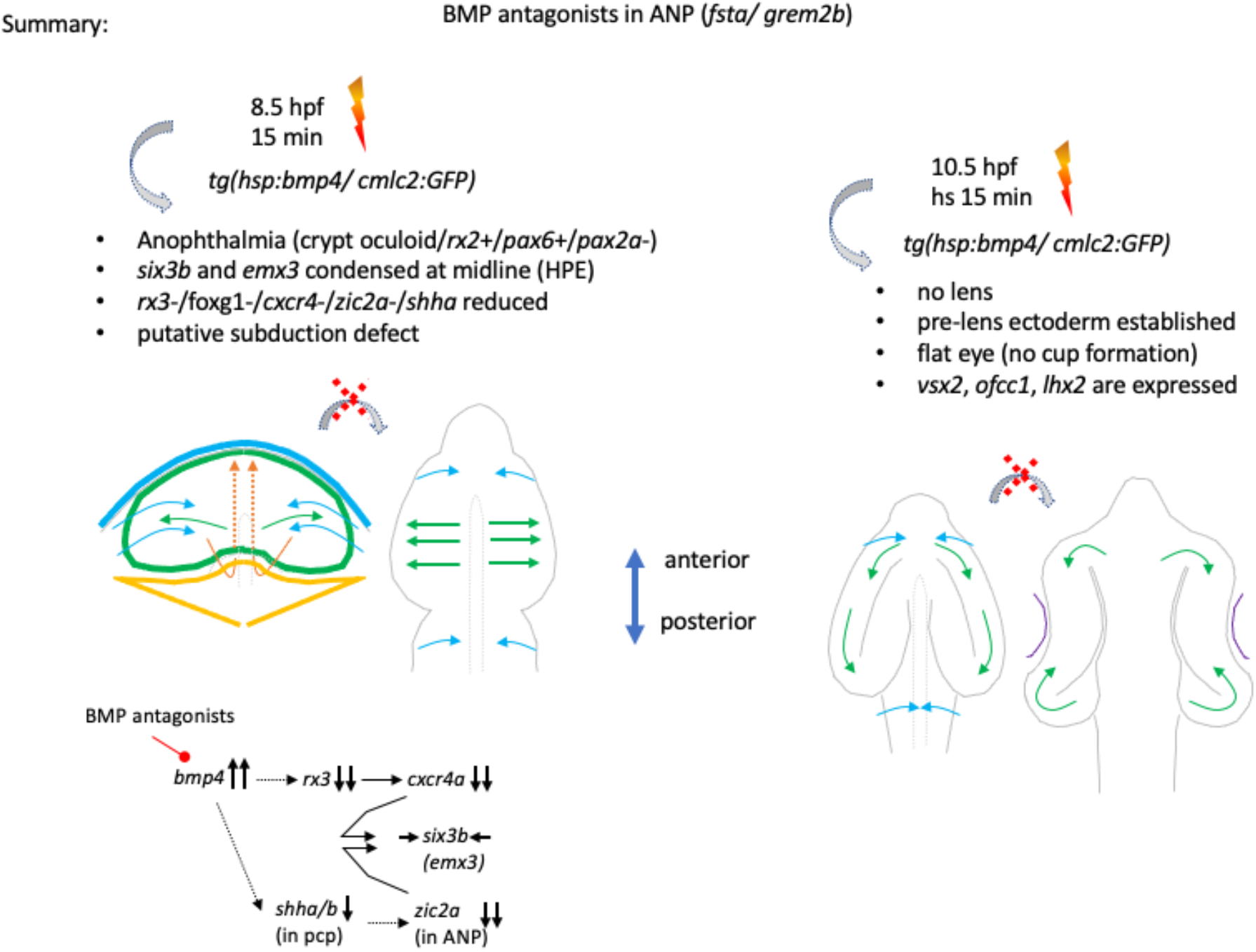
**Summary of major findings. Experimental induction of bmp4 and saturation of bmp antagonists in the ANP results in two phenotypes, hampered forebrain cleavage (HPE) (left) and absence of optic cup formation (right). Taken together these phenotypes result in anophthalmia with a crypt-oculoid.**

One upstream regulator of *ofcc1* is *vsx2* (Gago-Rodrigues et al., 2015). However, the transcription factor *lhx2* was also found essential for the transformation of the optic vesicle into the optic cup (Porter et al., 1997).

We next addressed the expression of *ofcc1* (**Figure 7 M-P**). In controls *ofcc1* can be found among other domains in the optic cup (**Figure 7 M, N**) but expression could also be found in the region of the oculoid after induction of *bmp4* (**Figure 7 O, P**). Analysis of *vsx2* expression, a proven upstream regulator of ofcc1, showed expression in the optic cup in controls and expression in the oculoid in *bmp4* induced embryos (**Figure 7 Q-T, arrow**). These findings suggest that the process of basal constriction of the retinal progenitors is more likely affected by other means than by the level of ofcc1. Also, the expression of *lhx2* could be detected in the oculoid after *bmp4* induction, while in controls at this stage it was found in telencephalon, diencephalon and ventral optic cup (**Figure 7 U-X, arrows**). This is suggesting that it is less likely the key to the failure in optic cup formation after *bmp4* induction.

### Summary Discussion and Conclusion

Holoprosencephaly (HPE) is the most important and most frequent developmental forebrain disorder in humans. The incidence in live births is approximately 1/10000, yet the estimated incidence per conception is much higher, 0.4 % determined in abortions (Matsunaga and Shiota, 1977). Different forms of HPE have been described going along with severe phenotypes like cyclopia or anophthalmia or with milder phenotypes like coloboma.

We recently identified BMP antagonism to be essential for optic vesicle to optic cup transformation and found morphogenetic defects including coloboma resulting from experimental induction of a BMP ligand (Eckert et al., 2019; Heermann et al., 2015; Knickmeyer et al., 2018). We proposed that “morphogenetic coloboma” are more likely linked to HPE than coloboma due to a defect of pioneer cells within the fissure margins (Eckert et al., 2020) resulting in a fusion defect of the fissure margins (Chan et al., 2021).

In this study we set out to address the role of BMP antagonism during early forebrain and eye development. We found BMP antagonists, *grem2b* and *fsta*, expressed within and neighboring the ANP at 11 hpf and in order to saturate this domain, we induced *bmp4* at 8.5 hpf. We found a severe form of HPE associated with anophthalmia resulting from the *bmp4* induction. Analyses of the ANP revealed a splitting defect of the eye field and the prospective telencephalon. We found marker for the future telencephalon and the eye field, *emx3* and *six3b*, condensed to the midline. The proper interaction of the prechordal plate mesoderm with the developing prosencephalon, the ANP at this stage, is likely the most important step for proper cleavage of the telencephalic field and the eye field (Li et al., 1997) and most HPE related genes and also non-genic risk factors can be linked to this step (Roessler et al., 2018). Shh is the essential factor secreted from the prechordal plate (Sagai et al., 2019) and in turn is directing correct forebrain development. An upstream regulator for prechordal plate development and thus also of Shh signaling in this domain was found with ZIC2 (Warr et al., 2008).

The HPE phenotype in *Zic2* mutants in mouse was explained by a defect during mid-gastrulation, hampering the regular development of the prechordal plate (Warr et al., 2008). We found that *zic2a* expression was ceased in the ANP domain after induction of *bmp4* (this study). In our paradigm we induce *bmp4* at 8.5 hpf, a timepoint later than mid-gastrulation (Kimmel et al., 1995). We, however, also found a mild and variable decrease in expression of *shha/b* in the anterior region, the prechordal plate. We cannot exclude that the induction of *bmp4* affected the expression of *zic2a* and *shha/b* independently. We can also not be sure whether the lack of *zic2a* was functionally relevant for inducing HPE (this study) or whether the reduced level of *shha/b* in the prechordal plate was the major factor in our paradigm.

It was shown before that the expression of *six3b* was depending on Hh signaling (Sanek et al., 2009). The expression of *six3b* was not lost after *bmp4* induction, but was condensed to the midline (this study) suggesting that at least the level of Shh was sufficiently high for *six3b* induction. Zic2 was also found to act downstream of Shh to affect *six3* expression in the ANP in zebrafish (Sanek et al., 2009). This effect was, however, considerably later, at mid-somitogenesis, and had an major influence on the development of the prethalamus (Sanek et al., 2009).

Based on morphology and the expression of genes marking e.g. precursors of the different ANP domains our data suggest that the subduction movement, which in wildtypes is the driver for eye field splitting (England et al., 2006) is hampered after induction of *bmp4*. It will be interesting in future analyses to address further the link between the *bmp4* induction, the regulation of *shha/b* and *zic2a* and the subduction movement of the hypothalamic domain.

Notably, in mouse, the compound knock-out of two BMP antagonists, chordin and noggin, resulted in severe forebrain defects presenting cases of HPE with cyclopia but also more severe cases with aprosencephaly (Bachiller et al., 2000). The onset of chordin expression was found in the primitive streak, subsequently in the node and the axial mesendoderm and at midgastrula chordin and noggin expression were found overlapping (Bachiller et al., 2000). Here, also the expression of shh was negatively affected (Bachiller et al., 2000). Even though the phenotypes of the compound mouse mutants and ours were both showing severe HPE, there were also distinct differences. We found anophthalmia and crypt-oculoids. It was not reported whether in the cases of the compound mutants, which resulted in aprosencephaly, *rx2* and *pax6* expressing domains could be found in the remaining tissue of the otherwise lost/reduced forebrain. Milder HPE forms in the compound mutants showed cyclopia, while we observed anophthalmia. We asked what makes the difference between these two phenotypes.

We found that *rx3* and *foxg1* expression was lost from the eye field and the future telencephalon respectively. The loss of *foxg1* is well explaining the growth defects of the telencephalon, which we saw at 24 hpf. Astonishingly, however, was the finding of a ceased *rx3* expression, because at 24 hpf we found a condensed domain of *rx2* and *pax6a* positive tissue in the dysmorphic forebrain, the crypt-oculoid. In the zebrafish *rx3* mutants *chokh*, especially *rx2* expression was shown to be depending on *rx3* (Loosli et al., 2003) and our own analysis of *rx3* crispants (this study) are well in line with this. In the medaka *rx3* mutants *eyeless* (*el*), however, *rx2* expression could be detected (Loosli et al., 2001). This difference may be due to the species, but also the nature of the mutation could play a role. In the zebrafish *chokh* mutant a point mutation (s399) results in a premature stop and thus in a truncated *rx3*. In medaka *eyeless*, on the other hand, an intronic insertion results in a transcriptional repression, which is also temperature sensitive. In our *bmp4* induction paradigm we do not mutate the *rx3* locus, but rather also regulate the expression of *rx3*. It is conceivable that *rx3* is expressed at a level which is not detectable with WMISH but sufficient to induce *rx2* expression in our paradigm and maybe also in the medaka eyeless embryos. In both, the level of *rx3* was, yet, not sufficient to allow for eye field splitting and optic vesicle out-pocketing (Rembold et al., 2006, this study). Irrespectively of the exact level of *rx3* expression/repression, we detected a dramatic reduction of *cxcr4a* expression within the eye field. *Cxcr4* was show to act downstream of *rx3* and together with *sdf1b* from the surface mesendoderm to facilitate the evagination of the eye field (Bielen and Houart, 2012). The “loss” of *rx3* and subsequently of *cxcr4a* in our *bmp4* induction paradigm, thus, very likely explains the lack of optic vesicle out-pocketing (this study). Yet, this did not explain why not a single cyclopic eye was formed.

In order for an optic vesicle to transform into a cup an interaction with the pre-lens ectoderm is needed (Hyer et al., 2003). A delayed but strong induction of *bmp4* was preventing lens formation and even dispersed lens tissue was not found. This could be used as an explanation for the resulting optic cup formation defect. After an early induction of *bmp4* we also did not observe a morphological lens, however, we found dispersed *cryaa* expressing tissue in a central anterior region, where in a cyclopic individuum the lens would be found. This indicated that the pre-lens ectoderm was existent, yet the morphogenesis of an optic cup was severely disturbed. We thus speculated that another reason must be causing anophthalmia instead of cyclopia after early *bmp4* induction. The motor for cup formation is the individual basal constriction, facilitated by opo/ofcc1 (Bogdanović et al., 2012; Martinez-Morales et al., 2009b) and our flattened optic vesicles reminded us of the *opo* mutant in medaka (Martinez-Morales et al., 2009b). *Vsx2* was shown to regulate expression of ofcc1/opo (Gago-Rodrigues et al., 2015). Since we did not detect a loss of *vsx2* or *ofcc1* expression after *bmp4* induction but found a flat optic vesicle, we deduced that the process of basal constriction must be hampered by another mechanism, which is to be identified in future analyses. We further speculated that the transcription factor *lhx2* could also be involved, since in lhx2 mouse mutants, the optic vesicle to optic cup transformation fails (Porter et al., 1997). But also the expression of *lhx2* was not lost after *bmp4* induction. Taken together our data suggest, that the failure of the initiation of optic cup formation, on top of the HPE observed after *bmp4* induction, is the reason for anophthalmia, even though a crypt-oculoid is existent. It will be interesting to address in future analyses, how *bmp4* induction affects the process of basal constriction, hampering optic cup formation.

## Materials and Methods

### Zebrafish care

Zebrafish were kept in accordance with local animal welfare law and with the permit 35-9185.64/1.1 from Regierungspräsidium Freiburg. Fish were maintained in a constant recirculating system at 28°C on a 12h light: 12h dark cycle. The following transgenic lines were used: *tg(hsp70l:bmp4, myl7:eGFP)*(Knickmeyer *et al*., *2018) tg(Ola*.*rx2:bmp4, myl7:eGFP)* (He*ermann et al*., *2015)*. Zebrafish embryos were grown in petri dishes in zebrafish medium, consisting of 0.3g/l sea salt in deionized water. If melanin-based pigmentation needed to be inhibited for downstream applications, embryos were grown in 0.2mM phenylthiourea.

### Heat shock procedures

For induction of heat-shock inducible transgenes, embryos were transferred to 1.5ml reaction tubes and incubated at 37°C in a heating block (Eppendorf Thermomixer). The onset and the duration of the respective heat shocks varied details are given in the results.

### Laser Scanning confocal Microscopy

Confocal images were recorded with an inverted TCS SP8 microscope (Leica). Embryos were embedded in 1% low-melting agarose (Roth) in glass-bottom dishes (MatTek). Live embryos were anaesthetized with MS-222 (Tricaine, Sigma-Aldrich) for imaging. Image stacks were recorded with a z-spacing of 3 µm, unless specified otherwise.

### Image Processing

Images from microscopy were edited for presentation using ImageJ (Fiji) software (Schindelin et al., 2012).

### In situ hybridization

Whole-mount ISH was performed according to (Quiring et al., 2004). For probes used please visit the Supplemental Data. Probes for cryaa and ofcc1 were designed cloning free (Hua et al., 2018).

### CRISPR/Cas9 F0 analysis (Crispants)

Embryos in 1-cell stage were microinjected with 1µM Cas9 protein (Alt-R S.p. Cas9 Nuclease V3, 1081059, Integrated DNA Technologies) and 1µg/µl sgRNA mix as described (Wu et al., 2018). sgRNAs were designed using CCTop (http://crispr.cos.uni-heidelberg.de) (Stemmer et al., 2015). For sequences please visit the Supplemental Data.

## Acknowledgements

We thank all members of the Heermann lab for insightful discussions and Ute Baur for great technical assistance. We want to thank Eleni Roussa and Klaus Unsicker for support of the Heermann lab.

