## Supplemental Data for "Early eye and forebrain development are facilitated by Bone Morphogenetic Protein antagonism"

Supplemental Data 1: Probes used for ISH

Alcamb: F: 5'-CACCCTCTCGCTACAACTTC-3'; R: 5'-CTTCTTGCTCTCCTCTGCTG-3'

Cryaa F: 5'-CCAACACCCTTGGTTCAGAC-3'; R: 5'-GTAACAGGGATGGTGCGATCT-3'

Cxcr4a F: 5'-TGGGTTGCCAGAAGAAATCC-3'; R: 5'-CCAGAAAGGCGTACAGGATC-3'

Emx3 F 5'-GAAGTGCTTCACGATTGAATC-3'; R 5'-TGAAATGACGTCAATGTCCTC-3'

Fgf8a F: 5'- GACTCATACCTTCACGGTTGAG-3'; R: 5'- TGCGTTTAGTCCGTCTGTTG-3'

Foxg1a F: 5'- ATGTTGGATATGGGAGAAAG -3'; R: 5'- AAGAAATAACTGGTCTGACC -3'

Fsta see Knickmeyer et al. 2018

Grem2b see Knickmeyer et al. 2018

Her13 F: 5'- CCACGCTGCTGAACTTAGAAA-3'; R: 5'- TCATCCAGGTCAGAGCAGAGA-3'

Ofcc1 F: 5'- GATGCTGCCAAGCTCTACTGG-3'; R: 5'- TTCATCCTCTCGTGTTGCTCAT-3'

Pax2a see Eckert et al. 2019

Pax6a F: 5'- AGATGGTTGCCAACAGTCAG -3'; R: 5'- GGGACATGTCTGGTTCACTG -3'

Rx2 F: 5'- GCCTCTCCACAGAAAGCTAC -3'; R: 5'- CGATACTAGAACTGCGGTCG -3'

Rx3 F: 5'- ATGAGGCTTGTTGGATCTCAG -3'; R: 5'- ATGAGGCTTGTTGGATCTCAG -3'

Shha see Knickmeyer et al. 2021

Shhb see Knickmeyer et al. 2021

Six3b F: 5'- TTTGGTCGTTGCCCGTAGCACC -3'; R: 5'- CATCGAAATCAGAGTCACTGTC -3'

Vsx2 see Eckert et al. 2020

Zic2a F: 5'- ATTAAGCAAGAGCTCATCTG -3'; R: 5'- AACTGTGGACCGCTGAGGAAG -3'

Supplemental Data 2: Sequences for CRISPR/Cas9

*rx3* T1: CCCGGCGTTTCCATATGGAT

*rx3* T2: TGAACGTGGTTCGGTTCCGC

*rx3* T3: CTTCGAGAAGTCGCACTATC

*rx3* T4: GAGATGGGGCCGGTCAACCA
